# Analysis host-recognition mechanism of staphylococcal kayvirus ɸSA039 reveals a novel strategy that protects *Staphylococcus aureus* against infection by *Staphylococcus pseudintermedius Siphoviridae* phages

**DOI:** 10.1101/632646

**Authors:** Aa Haeruman Azam, Kenji Kadoi, Kazuhiko Miyanaga, Masaru Usui, Yutaka Tamura, Longzhu Cui, Yasunori Tanji

## Abstract

Following the emergence of antibiotic resistant bacteria such as methicillin-resistant *Staphylococcus aureus* (MRSA) and methicillin-resistant *Staphylococcus pseudintermedius* (MRSP), phage therapy has attracted significant attention as an alternative to antibiotic treatment. Bacteriophages belonging to kayvirus (previously known as Twort-like phages) have broad host range and are strictly lytic in *Staphylococcus* spp. Previous work revealed that kayvirus ɸSA039 has a host-recognition mechanism distinct from those of other known kayviruses: most of kayviruses use the backbone of wall teichoic acid (WTA) as their receptor; by contrast, ɸSA039 uses the β-N-acetylglucosamine (β-GlcNAc) residue in WTA. In this study, we found that ɸSA039 could switch its receptor to be able to infect *S. aureus* lacking the β-GlcNAc residue by acquiring a spontaneous mutation in open reading frame (ORF) 100 and ORF102. Moreover, ɸSA039 could infect *S. pseudintermedius*, which has a different WTA structure than *S. aureus*. By comparison with newly isolated *S. pseudintermedius*–specific phage (SP phages), we determined that glycosylation in WTA of *S. pseudintermedius* is essential for adsorption of SP phages, but not ɸSA039. Finally, we describe a novel strategy of *S. aureus* which protects the bacteria from infection of SP phages. Notably, glycosylation of ribitol phosphate (RboP) WTA by TarM or/and TarS prevents infection of *S. aureus* by SP phages. These findings could help to establish a new strategy for treatment of *S. aureus* and *S. pseudintermedius* infection, as well as provide valuable insights into the biology of phage–host interactions.

## Introduction

*Staphylococcus* is a Gram-positive bacterium that causes many kinds of infections. Two representative species, *S. aureus* and *S. pseudintermedius*, are coagulase-positive bacteria that are notoriously pathogenic in humans and animals (Kloos and Bannerman 1994; Pompilio et al. 2015). *S. aureus* is a commensal found on the skin and mucosae of humans, whereas *S. pseudintermedius* commonly inhabits dog skin. Both are common bacterial pathogens associated with chronic and recurrent skin infections that require long-term systemic antimicrobial therapy. Infection by *S. aureus* and *S. pseudintermedius* is becoming problematic due to the emergence of antibiotic-resistant strains, including methicillin-resistant (MRSA and MRSP) and vancomycin-resistant strains (VRSA) (Enright et al. 2002; Sakoulas et al. 2005). Virulent bacteriophages that can kill a wide range of *S. aureus* hosts represent promising alternatives to conventional antibiotic treatment (Alves et al. 2014; Iwano et al. 2018; Azam and Tanji 2019a). The success of phage infection depends on its host specificity, which is often determined by the interaction between a phage receptor-binding protein (RBP) and its cognate receptor on the surface of the host cell (Hyman and Abedon 2010).

Staphylococcal kayviruses (previously known as Twort-like phages) have broad host range and high lytic ability, making them suitable candidates for phage therapy (Łobocka et al. 2012). Most phages belonging to this group use the backbone of wall teichoic acid (WTA), the most abundant molecule in the cell wall of *Staphylococcus*, as their receptor (Xia et al. 2011). However, previous work showed that kayvirus ɸSA039 uses the β-GlcNAc moiety in WTA of *S. aureus* SA003, a unique feature within the group (Azam et al. 2018).

The broad host range of kayviruses includes coagulase-negative staphylococci (CoNS) (Cui et al. 2012; Lobocka et al. 2012; Iwano et al. 2018). WTA in *S. aureus* generally consists of repetitive 1,5-ribitol-phosphate (RboP) modified with a GlcNAc residue and D-alanine. The GlcNAc moieties are transferred onto WTA by the α-GlcNAc transferase TarM and the β-GlcNAc transferase TarS (Xia et al. 2010; Brown et al. 2012). By contrast, WTA of *S. pseudintermedius* and CoNS has glycerol-phosphate (GroP) as the backbone and various glycoepitopes (GlcNAc, GalNAc, or Glc) (Endl et al. 1983; Winstel et al. 2014).

Staphylococcal phages that use glycoepitopes in the WTA as their receptors, e.g., phages from families *Siphoviridae* and *Podoviridae*, target either the RboP or GroP type of WTA. For example, *Siphoviridae* ɸ11 only recognizes GlcNAc of RboP WTA, whereas ɸ187 only recognizes the glycoepitope of GroP WTA (Winstel et al. 2014). Therefore, the unique feature of ɸSA039, a kayvirus that requires the β-GlcNAc residue in the WTA of SA003 (RboP WTA), raises the question of whether this phage can also recognize other types of WTA from different *Staphylococcus* species. In this study, we evaluated the host range of ɸSA039 and its potential for phage therapy toward various strains of *S. pseudintermedius*, including MRSP. In addition, we also screened *S. pseudintermedius*–specific phages, evaluated them as potential antimicrobial agents, and compared them with ɸSA039.

In addition, a previous study showed that kayviruses such as ɸSA012 and ɸK have at least two RBPs (RBP1 and RBP2), which are responsible for these phages’ broad host range (O’Flaherty et al. 2004; Takeuchi et al. 2016). Mutant ɸSA012 has a modified RBP2 (ORF103) that enables the phage to infect mutant *S. aureus* SA003R38, which has an altered WTA and overproduces capsular polysaccharide (Takeuchi et al. 2016; Osada et al. 2017; Azam et al. 2018), indicating that mutated RBP2 may allow the phage to counter-adapt to resistant hosts by using an alternative component as a novel receptor (Takeuchi et al. 2016). Like ɸSA012, ɸSA039 also has two RBPs, RBP1 (ORF100) and RBP2 (ORF102) (Azam et al. 2018). Hence, in light of the unique infection strategy of ɸSA039, we also investigated the ability of the phage to counter-adapt to phage-resistant host and analyzed the underlying mechanism.

## Material and Methods

### Bacterial strains, bacteriophages, and plasmids

Bacteria, phages, and plasmids used in this study are listed in Table 1. *S. aureus* RN4220 was used with the permission of Professor Richard P. Novick (Skirball Institute of Biomolecular Medicine, New York, NY, USA). *S. aureus* SA003 was isolated from milk of a mastitic cow (Synnott et al. 2009). The *S. aureus* virulent phages ɸSA039 and ɸSA012 were isolated from sewage in Japan (Synnott et al. 2009). All *S. pseudintermedius* isolates were isolated from the skin of dogs with canine pyoderma. Nine coagulase-negative *Staphylococcus* (CoNS) were isolated from a patient at Jichi Medical University hospital, and species identification was performed based on the 16S rRNA gene. *S. aureus* SA003, *S. pseudintermedius* SP015, *S. pseudintermedius* SP070, and *S. pseudintermedius* SP079 were deposited in the culture collection of NITE Biological Research Center, Kisarazu, Japan, under accession numbers NBRC110650, NBRC113855, NBRC113857 and NBRC113858, respectively. All phages used in this study are deposited in the corresponding author’s institution and distributed to other researcher by request. All primers used in this study are listed in Supplemental Table S1.

**Table 1.**
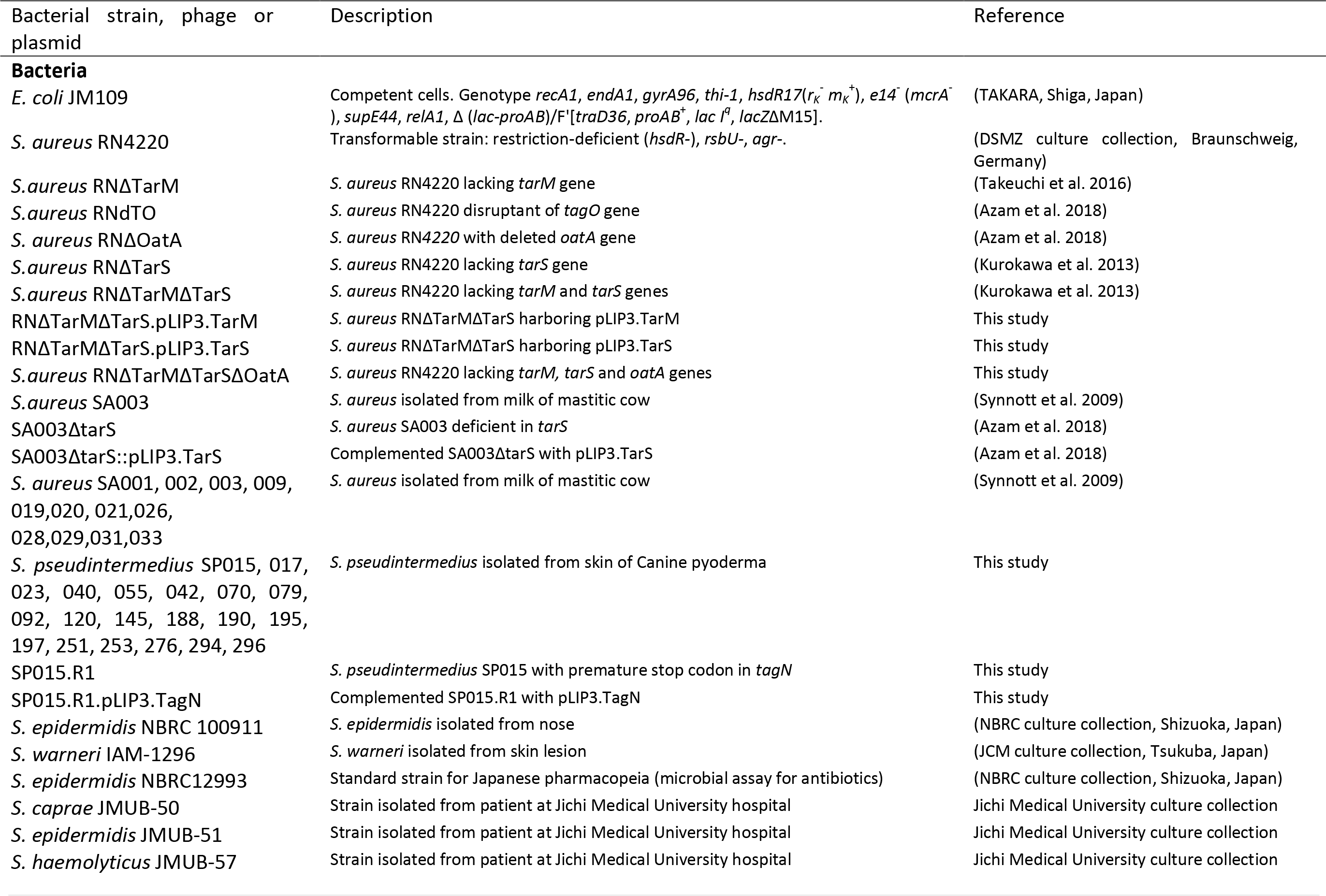
Bacterial strains, phages and plasmids used in this study

### Isolation of S. pseudintermedius

*Staphylococcus* was isolated based on standard culture and biochemical tests on site. Screening of *S. pseudintermedius* was performed using the API®/ID32 kit (Sysmex-bioMérieux, Tokyo, Japan). To distinguish *S. pseudintermedius* and *S. intermedius*, restriction fragment length polymorphism (RFLP) analysis was performed on the *kat* gene (Blaiotta et al. 2010). *S. pseudintermedius* strains were distinguished by PCR amplification of the hypervariable X region of the protein A gene (*spa*) (Moodley et al. 2009). Multilocus sequence typing (MLST) was performed following Bannoehr’s method, which targets five genes (16SrRNA, *tuf*, *cpn60*, *pta*, and *agrD*) (Bannoehr et al. 2007; Bannoehr et al. 2009).

### Bacteriophage isolation and purification

Phage ɸDP001, which is capable of lysing *S. pseudintermedius*, was isolated from dog saliva. Saliva was collected using sterile cotton swabs, which were soaked in 1 ml SM buffer (100 mM NaCl, 8 mM MgSO_4_, 50 mM Tris-HCl [pH 7.5], 0.01% gelatin), and incubated at 4°C for at least 1 h. After centrifugation (9730 *g*, 1 min), 100 μl supernatant was mixed with 100 μl overnight culture of *S. pseudintermedius* SP015 in 3 ml of 0.5% top agar, plated on LB agar, and incubated at 37°C overnight. ɸSP120, ɸSP197, and ɸSP276 were isolated by mitomycin induction; Mitomycin was added into *S. epidermidis* culture (OD = 0.5) at a final concentration of 0.5 μg/ml and incubated for 1 h. Phages were isolated from the supernatant after centrifugation (9730 *g*, 3 minutes). Propagation and purification of the phages was performed as previously described (Synnott et al. 2009).

### Plaque assay and efficiency of plating (EOP)

Plaque assay was performed by mixing 100 μl of phage (10^4^ PFU/ml) with 100 μl overnight culture of bacteria in 3 ml of 0.5% top agar, plated on LB agar, and incubated at 37°C overnight. Experiments were conducted in triplicate. The EOP value was measured as a percentage of the number of observed plaques on the tested bacteria divided by the number of plaques observed on the wild-type strain.

### Spot test

Two microliters of phage lysate at a titer of 10^8^ plaque forming units (PFU) per ml (10^5^ PFU) was dropped onto a lawn of bacteria and incubated overnight at 37°C. Experiments were conducted in triplicate.

### Adsorption assay

The adsorption efficiency of phages on *S. aureus* strains was measured by titrating the presence of free phages in the supernatant after 20 minutes of cell–phage contact following previous study (Azam et al. 2018). Free phage was collected by centrifugation (9730 × *g*, 1 min) and titrated using SA003 for ɸSA039 and SP070 for SP phages.

### Isolation and characterization of spontaneous mutant of ɸSA039

Co-culture experiments were performed with *S. aureus* SA003 and ɸSA039. First, 4.5 ml of LB broth was inoculated with 1% overnight culture of SA003 and cultured until early exponential phase (OD_660_=0.1) in a TVS062CA BioPhotorecorder (Advantec, Tokyo, Japan). Then, phage ɸSA039 was added at a multiplicity of infection (MOI) = 1. The mixture was cultured at 37°C with shaking at 40 rpm. After 2 days, the culture was transferred to 4.5 ml of fresh LB medium (1% dilution) and cultured under the same condition. Co-cultures were repeated until a spontaneous mutant of ɸSA039 that could infect TarS-null *S. aureus* was isolated. Two spontaneous mutants were isolated from the co-culture and were designated ɸM1 and ɸM2.

### Checking for mutations in genes encoding tail and baseplate proteins

Genes encoding tail and baseplate proteins (ORF95–ORF102) in spontaneous mutant phages were amplified by plaque PCR using DirectAmpPCR (TAKARA, Shiga, Japan) and analyzed by Sanger sequencing. Phage plaques were touched with a toothpick and mixed into the PCR mixtures.

### Generation and characterization of chimeric phage

Chimeric phage was generated by homologous recombination using plasmid pNL9164 (Sigma-Aldrich, MO, USA) in *S. aureus* SA003. Mutated DNA fragments of ORF100 and ORF102 were amplified from spontaneous mutant phages by PCR using KOD-plus Neo enzyme (Toyobo, Shiga, Japan). PCR fragments and the plasmid were digested with appropriate restriction enzymes. The DNA fragment was inserted into plasmid pNL9164 using T4 DNA Ligase (New England BioLabs, Ipswich, MA, USA). The constructed plasmid was cloned into *Escherichia coli* JM109 competent cells (TAKARA, Shiga, Japan) and pre-introduced into restriction-deficient *S. aureus* RN4220 before being transformed into *S. aureus* SA003.

To perform homologous recombination, transformant SA003 harboring plasmid with homologous region was infected with phage (MOI = 1). The mixture was incubated at 37°C overnight. Recombinant phages were isolated from the supernatant fraction of the mixture after centrifugation (8000 × g, 3 minutes). SA003ΔTarS was used to screen recombinant phages from homologous recombination of ORF100, and RN4220 was used to screen recombinant phages from homologous recombination of ORF102. Mutated ORF100 from spontaneous mutant phage ɸM1 was introduced into wild-type ɸSA039, yielding chimeric phage ɸrM1/r-100. Mutated ORF102 of ɸM1 was introduced into ɸrM1/r-100, yielding chimeric phage ɸrM1/r-100&102.

### Deletion of *oatA* gene in RN4220ΔtarMΔtarS

The gene was deleted by pCasSA plasmid with clustered regularly interspaced short palindromic repeats (CRISPR)-Cas9 system (Chen et al. 2017). Plasmid construction was performed as previously described (Chen et al. 2017; Azam et al. 2018). Spacers were manually selected by searching the protospacer adjacent motif (PAM) region. Two oligos were designed as single-stranded DNA for each spacer, and double-stranded spacer was generated by phosphorylation with T4 PNK (New England BioLabs, Ipswich, MA, USA) and annealed at 95°C for three minutes. Plasmid pCasSA was digested with *Bsa*I. The double-stranded spacer and digested pCasSA were ligated with T4 DNA ligase (New England BioLabs, Ipswich, MA, USA). Editing template was amplified by splicing by overlap extension (SOE) PCR using the region flanking the target gene as the DNA template. The resultant plasmid was cloned into *Escherichia coli* JM109 competent cells (TAKARA, Shiga, Japan) and transformed into *S. aureus* RN4220ΔtarMΔtarS. Complementation of deletion mutants was performed using plasmid pLI50 (purchased from Addgene [Cambridge, MA]) under the control of the P3 promoter (pLIP3) (Jeong et al. 2011; Takeuchi et al. 2016). The wild-type allele from RN4220 was used as the insert.

### Selection of phage-resistant mutants of *S. pseudintermedius* SP015

Plaque assays were performed using SP015 and ɸSA039 at an MOI = 10 and incubated at 37°C overnight. Colonies (resistant mutants) were purified, inoculated onto LB plates, and incubated at 37°C overnight. The phage resistance of isolated mutants was determined by spot test and adsorption assay. Phage-resistant SP015-R1 which showed ability to inhibit phage adsorption was further characterized.

### DNA extraction, sequencing, and bioinformatics

Phage genome was extracted using phage DNA isolation kit (NORGEN, ON, Canada). Bacterial genome was extracted using DNA extraction kit (Sigma-Aldrich, St. Louis, MO, USA). Whole-genome sequencing was conducted on the Illumina HiSeq platform with genome coverage (sequencing depth) of 100-fold. Genomes were assembled using velvet ver 1.2.10. ORFs were predicted and annotated using the RAST server (http://rast.nmpdr.org/). The presence of toxin or virulence genes in the phage genome was determined using PHASTER (PHAge Search Tool Enhanced Release) server (phaster.ca).

### Statistical analysis

Two-tailed Student’s *t-*test was used to determine statistical significance.

### Accession number (s)

Genome data of three SP phages (ɸSP120, ɸSP197, and ɸSP276) and *S. pseudintermedius* SP079 were submitted to the DNA Data Bank of Japan (DDBJ) database under accession numbers AP019560, AP019561, AP019562, and AP019372, respectively.

## Results

### ɸSA039 can recognize WTA of *Staphylococcus pseudintermedius* SP015

ɸSA039 exhibited at least a moderate ability to infect various strains of *S. pseudintermedius*, and 12 CoNS (Table 2). A close relative of ɸSA039, ɸSA012, also had a broad host range toward the *Staphylococcus* species we tested. In this study, we also isolated *S. pseudintermedius–*specific phages (SP phages) ɸDP001, ɸSP120, ɸSP197, and ɸSP276, and compared them with ɸSA012 and ɸSA039.

**Table 2.**
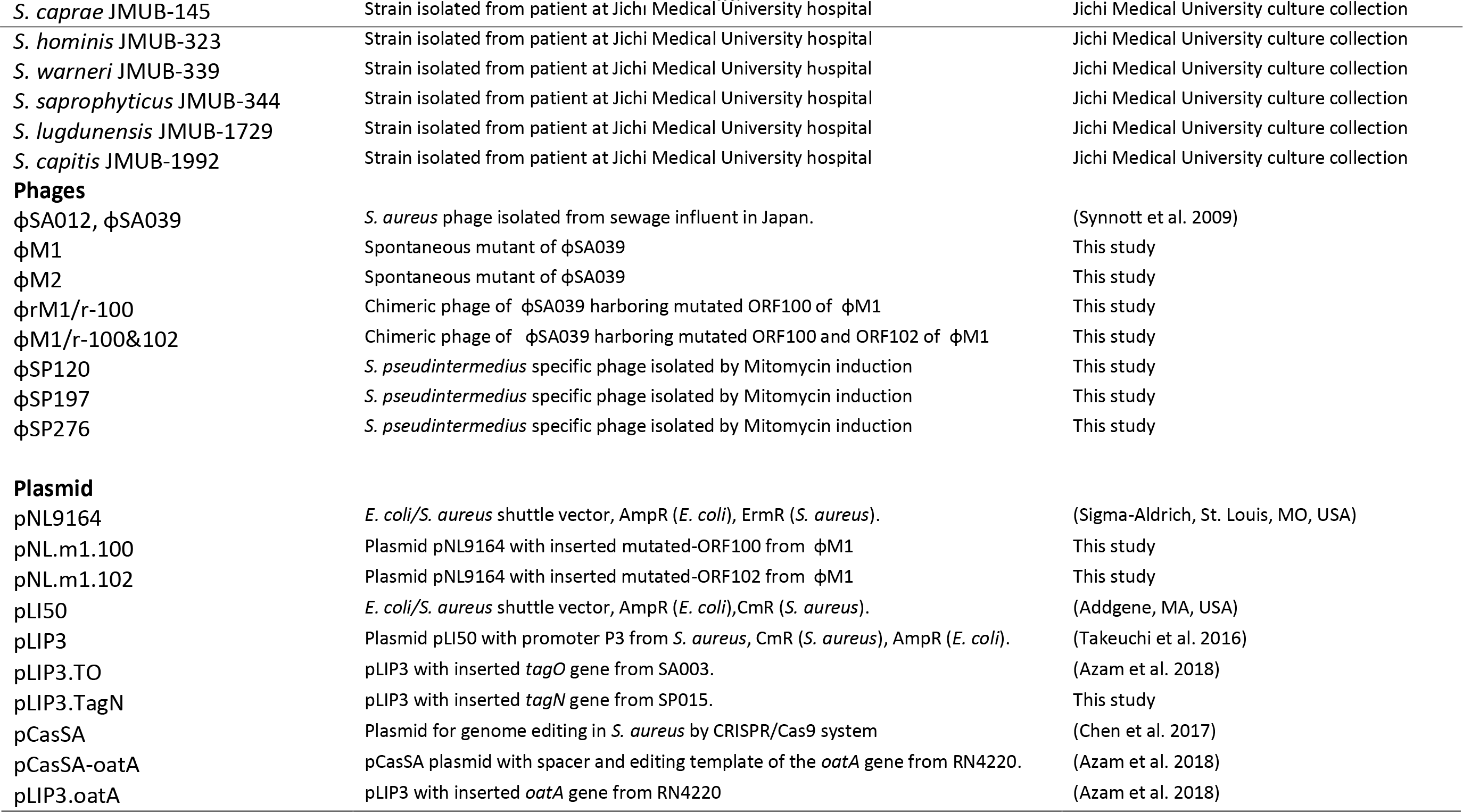
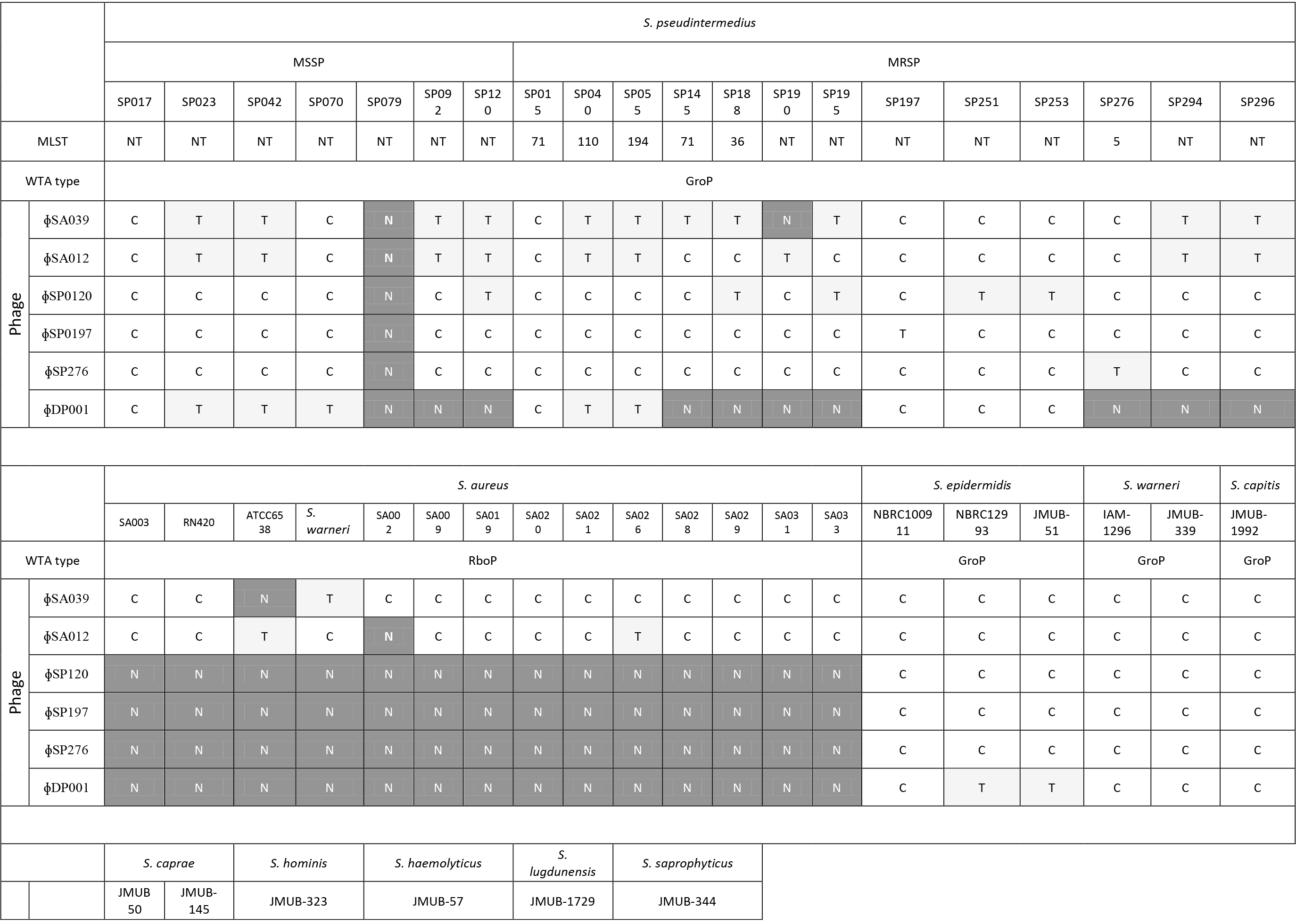
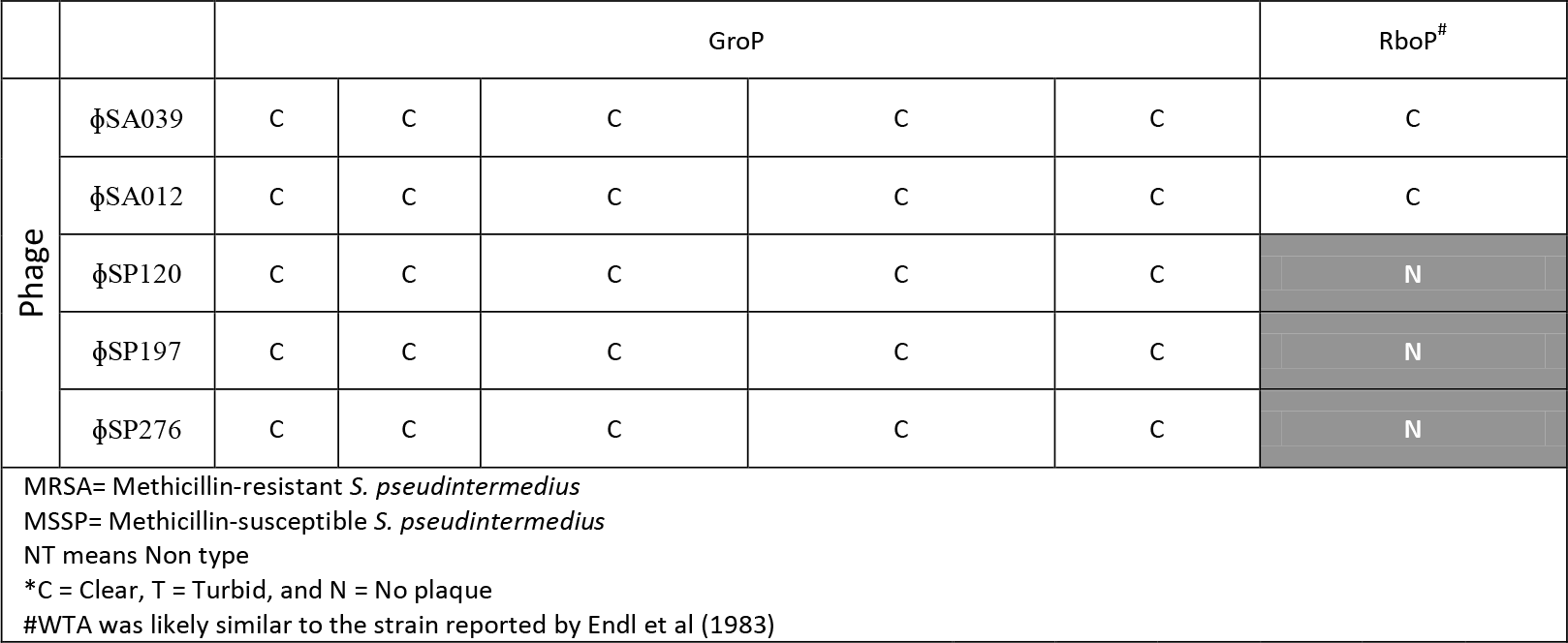
Spot test of phages toward *Staphylococcus aureus*, *S. pseudintermedius* and 12 CoNS

As shown in Table 2, in 20 isolates of *S. pseudintermedius*, ɸSA012 and ɸSA039 exhibited at least a moderate infection (turbid plaque): ɸSA012 infected 95% of isolates (19/20), whereas ɸSA039 infected 90% (18/20). ɸSA012 and ɸSA039 could infect 93% of *S. aureus* isolates (13/14) and the 12 CoNS. Unlike ɸSA012 and ɸSA039, no SP phages could infect the *S. aureus* strains used in this study. Regarding *S. pseudintermedius*, ɸDP001 infected 50% of isolates (10/20), whereas the other three SP phages infected 95% of isolates (19/20). *S. pseudintermedius* SP079 was the only strain that was resistant against all SP phages. Whole-genome analysis revealed that this bacterium encodes CRISPR-Cas9 system in its genome. DNA fragments of ɸSP120 and ɸSP197 were identified in the CRISPR regions of the two SP079 (Supplementary Table 1). Four SP phages formed at least turbid plaques in CoNS, with the exception of *Staphylococcus saprophyticus*.

We speculated that SP phages recognize receptors in *S. pseudintermedius* and CoNS isolates that are absent in *S. aureus*. Because most staphylococcal phages require WTA as their receptor (Azam and Tanji 2019b), we sought to determine whether our SP phages also require WTA. To this end, we generated WTA-free *Staphylococcus* by inhibiting WTA synthesis in the cell using tunicamycin (Zhu et al. 2018). ɸSA039, ɸSA012, and three SP phages (ɸDP001, ɸSP120, and ɸSP197) had no ability to infect WTA-free *S. pseudintermedius* and *S. epidermidis* (data not shown). The phages could not form a plaque and failed to adsorb onto WTA-free isolates. By contrast, ɸSP276 did not completely lose its infectivity toward WTA-free isolates, indicating that phage can use another component as a receptor. These findings indicated that all phages in this study utilize WTA in *S. pseudintermedius* and *S. epidermidis* as their receptor, but that the WTA of these bacteria is likely distinct from that of *S. aureus*.

### Accumulation of point mutations in ORF100 enables ɸSA039 to infect TarS-null *S. aureus*

ɸSA039 requires β-GlcNAc glycosylation of WTA by TarS (Azam et al. 2018). In this study, we found that ɸSA039 could generate spontaneous mutants capable of infecting TarS-null *S. aureus*. Mutants of ɸSA039 that could infect TarS-null *S. aureus* were obtained from the fifth batch of two co-cultures. One spontaneous mutant phage (ɸM1and ɸM2) was purified from each of co-culture and further characterized. In SA003ΔtarS, adsorption of ɸM1and ɸM2 was around 8-fold than that of wild-type ɸSA039 (Fig 1a). ɸM1and ɸM2 exhibited similar adsorption toward SA003ΔtarS. Because the phage tail fiber and baseplate region are thought to be involved in phages’ adsorption specificity, we amplified the genomic region encoding the tail and baseplate proteins (ORF94 until ORF102) using primers described in Supplemental Table S1. Spontaneous mutations were detected in ORF100 and ORF102. ɸM1 harbors three point mutations in ORF100 and one point mutation in ORF102, whereas ɸM2 only has three point mutations in ORF100. All mutations changed the amino acid sequence of the protein. The mutations in ORF100 were distributed among five locations (one near the N-terminus, two in the middle, and two near the C-terminus). A point mutation located at base 623 (TCT, S →TAT, Y) was detected in both mutant phages. Point mutations at bases 907 (GAT, D → TAT, N) and 1850 (ACG, T → AGG, R) were detected in ɸM1, whereas mutations at bases 1012 (GAA, E → AAA, K) and 1844 (ACA, T → AAA, K) were detected in ɸM2. The mutation in ORF102 was located in the C-terminus at base 1116 (AAA, K → ACA, T).

**Fig 1.**
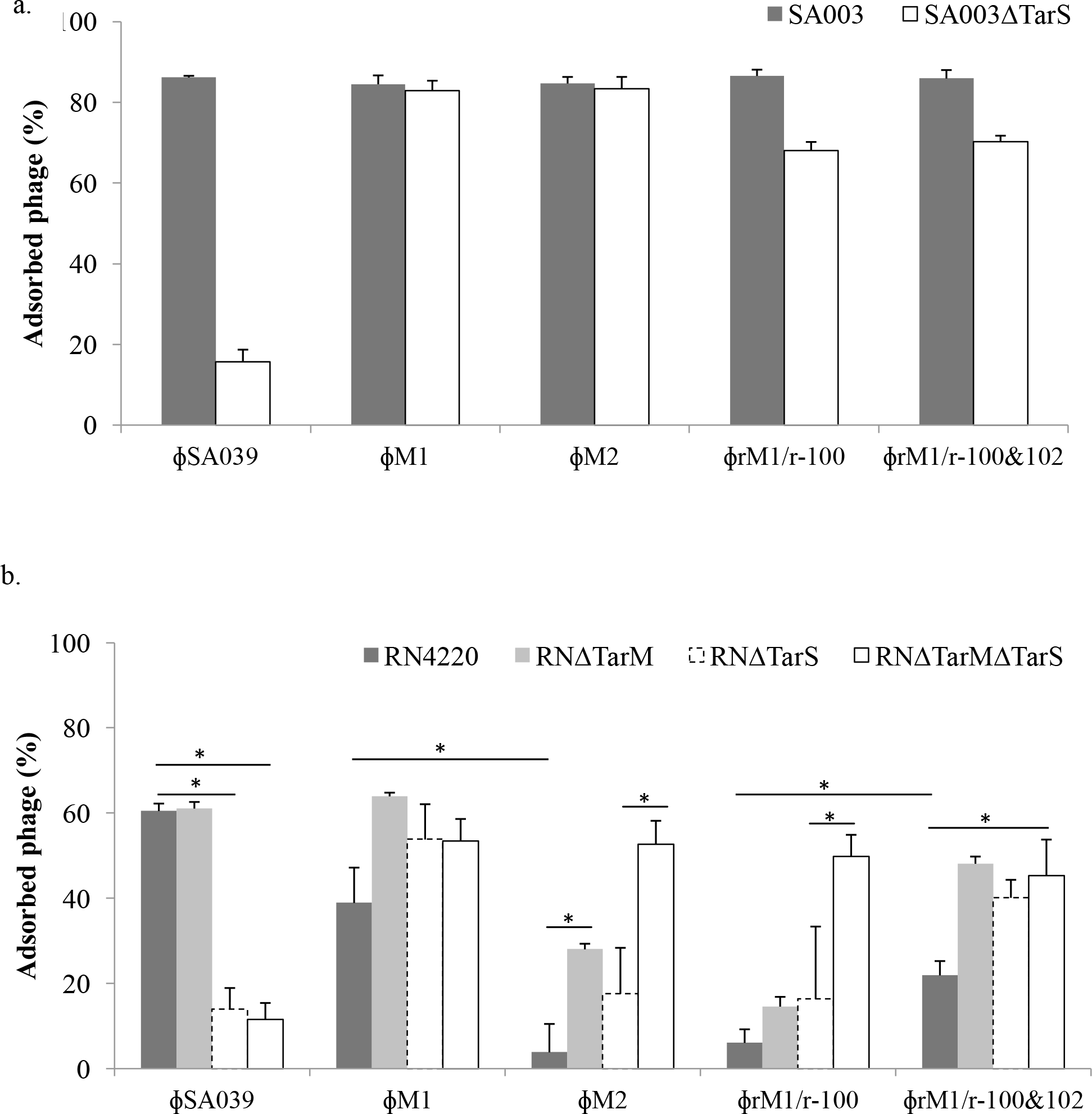
Adsorption of ɸSA039 and its derivative mutants (spontaneous mutant ɸM1 and ɸM2, recombinant phage harboring point mutations in ORF100 [ɸrM1/r-100], and recombinant phage harboring point mutations in ORF100 and ORF102 [ɸrM1/r-100&102]) onto (a) SA003 and *tarS* deletion mutant (SA003ΔTarS), (b) wild-type RN4220, *tarM* deletion mutant (RNΔTarM), *tarS* deletion mutant (RNΔTarS), and *tarM*/*tarS* double deletion mutant (RNΔTarMΔTarS). Statistical significance (*P*<0.05) is indicated with *.

Because all ɸSA039 mutants had spontaneous base changes in ORF100, we speculated that these mutations enabled the phages to infect SA003ΔtarS. We hypothesized that by introducing the point mutation in ORF100 into wild type ɸSA039, we should be able to construct chimeric ɸSA039 capable of infecting SA003ΔtarS. Hence, we introduced ORF100 of ɸM1 into wild-type ɸSA039 by homologous recombination, yielding the chimeric phage ɸrM1/r-100.

Spot tests revealed that ɸrM1/r-100 could infect SA003ΔtarS (data not shown). We then performed an adsorption assay to determine whether the ability of the recombinant phages to infect SA003ΔtarS was due to an effect of adsorption as a result of replacement of ORF100. Indeed, adsorption of ɸrM1/r-100 on SA003ΔtarS was significantly elevated relative to wild-type ɸSA039 (Fig. 1a).

### The α-GlcNAc residue in WTA blocks the infection of mutant ɸSA039 lacking a mutation in ORF102

*S. aureus* SA003 naturally lacks the gene encoding glycosyltransferase TarM (genome accession number: AP018376). Because certain *S. aureus* strains have *tarM* and *tarS* in their genome (Brown et al. 2012), in this study, we also used *S. aureus* RN4220, which has a complete set of WTAs (genome accession number: GCA_000212435.2), to characterize the mutant phage. As shown in Fig. 1b, wild-type ɸSA039 could not infect RN4220ΔtarS or RN4220ΔtarMΔtarS. However, mutant phages ɸM1 and ɸM2 exhibited completely different patterns of infectivity in the presence of *tarM*. Infection of ɸM2 was severely impaired toward RN4220, which has *tarM*. Only 3.86% of ɸM2 could adsorb onto RN4220, whereas 38.93% of ɸM1 could do so. Deletion of the *tarM* gene in RN4220 (RN4220ΔTarM) improved adsorption of ɸM1 and ɸM2 to 63.97% and 28.04%, respectively. A similar pattern was observed in RN4220ΔTarS. The presence of *tarM* in RN4220ΔTarS decreased the adsorption efficiency of both ɸM1 and ɸM2, and conversely, deletion of *tarM* in RN4220ΔTarS (RN4220ΔTarMΔTarS) improved the adsorption of both mutant phages. Overall, the mutations in ɸM1 and ɸM2 decreased the ability of the phages to adsorb onto WTA glycosylated by TarM; in particular, adsorption of ɸM2 was almost completely abolished.

As with ɸM2, the chimeric phage ɸrM1/r-100 was unable to adsorb onto RN4220 (6.06%), but was infectious toward RN4220ΔTarM (14.60% adsorbed phage). Based on this observation, we hypothesized that the absence of a point mutation in ORF102 in ɸM2 and ɸrM1/r-100 may make these phages unable to infect RN4220. To test this idea, we introduced a point mutation in ORF102 into ɸrM1/r-100 by homologous recombination, yielding the chimeric phage ɸrM1/r-100&102. Indeed, adsorption of the chimeric phage ɸrM1/r-100&102 improved three times higher than that of ɸrM1/r-100 (fig 1b).

### Whole-genome sequencing of SP phages

Because ɸSP120, ɸSP197, and ɸSP276 had a broad host range in *S. pseudintermedius*, we further investigated their potential as antimicrobial agents. To this end, we first sequenced the entire genomes of all three phages. In doing so, our primary goal was to investigate the presence of toxin, virulence, and antibiotic-resistance genes that could make them inappropriate for phage therapy.

The whole-genome sequencing analysis revealed that ɸSP120, ɸSP197, and ɸSP276 belong to *Siphoviridae*, with genome sizes of 40530, 41149, and 40711 bp, respectively. The GC content of ɸSP120, ɸSP197, and ɸSP276 was similar to that of *S. pseudintermedius*, but somewhat higher that of *S. aureus* (36%, 35%, and 36%, respectively). The genomes of the phages had a low degree of identity relative to one another. Their genomes were organized into six functional modules; lysogeny, DNA replication, packaging, head, tail, and lysis. The presence of integrase (CI) and repressor (Cro) genes in the genome indicate that these phages can undergo a lysogenic cycle (Xia and Wolz 2014). The integrases of the three phages were identical. Toxin, virulence, and antibiotic-resistance genes were absent from all three genomes, according to the PHASTER server.

Using the ViPTree online tool (https://www.genome.jp/viptree), we found that these phages are phylogenetically close to ɸ187 (Genome accession number: NC_007047). ɸ187 is a *Siphoviridae* phage isolated from the *S. aureus* ST395 lineage by mitomycin C induction (Asheshov and Jevons 1963). Other phages evolutionarily close to our SP phages were Stb27 (Genome accession number: NC_019914) and StB12 (Genome accession number: NC_020490), both of which were isolated from CoNS *Staphylococcus hominis* trough mitomycin C induction (Deghorain et al. 2012). The WTA structure of the *S. aureus* ST395 lineage and CoNS are similar to that of *S. pseudintermedius* (Winstel et al. 2014).

### WTA of *S. pseudintermedius* is different to that of *S. aureus*

Because SP phages cannot infect *S. aureus*, we speculated that the WTA of *S. pseudintermedius* is distinct from that of *S. aureus*. To date, the known WTAs of *Staphylococcus* bacteria are of at least two types, GroP and RboP (Brown et al. 2013; Winstel et al. 2014). Using primers targeting the *tagF* gene, which encodes the GroP polymerase of *S. pseudintermedius* ED99, we determined that all *S. pseudintermedius* in our study likely possess GroP type WTA. Most of the isolates had WTA clusters similar to that of ED99 (genome accession number: NC_017568). Genes encoding glycosyl transferases TarM and TarS were absent from *S. pseudintermedius* isolates. As with *S. pseudintermedius*, CoNS also have GroP WTA (Endl et al. 1983; Winstel et al. 2014). Therefore, our SP phages may specifically recognize GroP WTA, whereas ɸSA012 and ɸSA039 recognize both GroP and RboP WTA.

ɸ187 can utilize WTA of *S. aureus* PS187, which belongs to the ST395 lineage, but not that of the common *S. aureus* lineage (Winstel et al. 2014). Unlike other *S. aureus–*type strains, *S. aureus* ST395 has GroP WTA with α-GalNAc glycosylation mediated by TagN (Winstel et al. 2013, Winstel et al. 2014). In this study, we detected the presence of *tagN* in most *S. pseudintermedius* isolates. However, *S. pseudintermedius* SP070 and SP190 lacked *tagN* but had a gene encodes glycosyltransferase that absent from other isolates (unpublished data).

### SP phages can recognize non-glycosylated RboP WTA in *S. aureus*

WTA sugar modifications are highly variable in *Staphylococcus* species, and have been implicated in bacteriophage susceptibility and immunogenicity (Brown et al. 2013). The WTA of *S. aureus* contains α-GlcNAc and/or β-GlcNAc. In this study, we found that although no SP phages could infect *S. aureus* (Table 2), the absence of α-GlcNAc and β-GlcNAc in the WTA enabled the SP phages to infect this bacterium. SP phages could infect *S. aureus* RN4220ΔTarMΔTarS and SA003ΔTarS, both of which lack the GlcNAc modification in their WTA (Fig 2). Because SA003 naturally lacks TarM, SA003ΔTarS has a WTA similar to that of RN4220ΔTarMΔTarS. Although the observed plaques were slightly turbid, SP phages could adsorb onto RN4220ΔTarMΔTarS (Fig 2). Adsorption efficiencies of ɸSP120, ɸSP197, and ɸSP276 onto RN4220ΔTarMΔTarS were 40.34%, 32.69%, and 43.75%, respectively. Complementation of either TarM (α-GlcNAc) or TarS (β-GlcNAc) blocked infection by SP phages; however, the complete absence of WTA in RN4220dtagO also prevented infection. Thus, SP phages can utilize non-glycosylated WTA as their recognition site on *S. aureus* cells.

**Fig 2.**
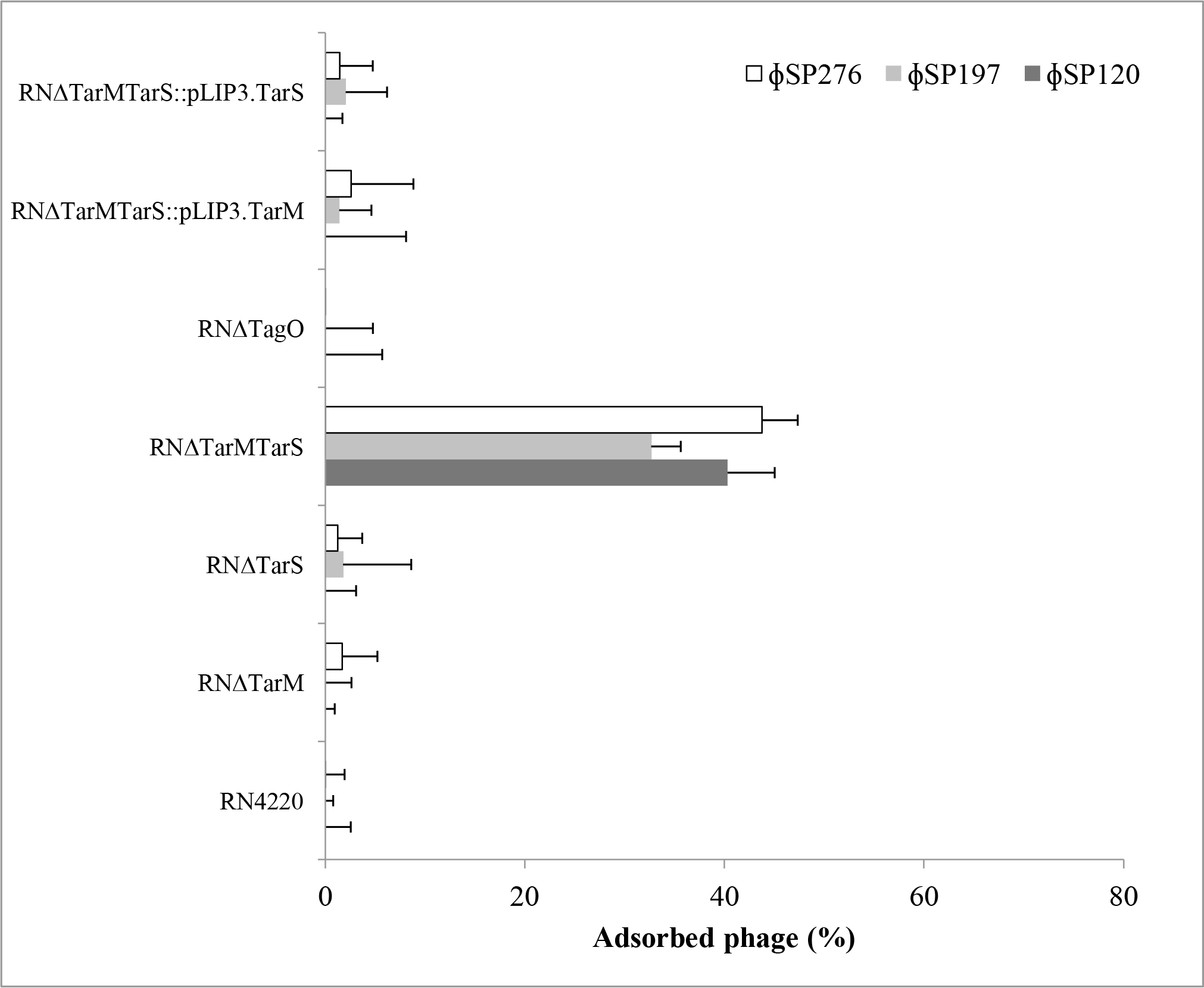
Infectivity of SP phages (ɸSP120, ɸSP197, and ɸSP276) toward RN4220 and its deletion mutants; *tarM* deletion mutant (RNΔTarM), *tarS* deletion mutant (RNΔTarS), and *tarM*/*tarS* double deletion mutant (RNΔTarMΔTarS), *tagO* deletion mutant (RNΔTagO), *tarM* complemented mutant RNΔTarMΔTarS::pLIP3.TarM, and *tarS* complemented mutant RNΔTarMΔTarS::pLIP3.TarS, as determined by adsorption assay.

6-O acetylation of muramic acid residues in peptidoglycan of *S. aureus* decreases the adsorption ability of staphylococcal *Siphoviridae* phages ɸ11 and ɸ52A (Li et al. 2016). To determine whether peptidoglycan acetylation is involved in SP phage adsorption onto RN4220 ΔTarMΔTarS, we generated a deletion mutant of *oatA* and used this strain as a host for the adsorption assay. We did not observe a change in the adsorption of SP phages relative to RN4220ΔTarMΔTarS (data not shown), suggesting that peptidoglycan acetylation is not essential for SP phage adsorption onto *S. aureus*.

### Glycoepitope of WTA in SP015 is essential for SP phages

Staphylococcal *Siphoviridae* phages require glycosylated WTA as their receptor (Xia et al. 2011). In RboP WTA, *Siphoviridae* phages recognize the GlcNAc residue, regardless of its stereochemistry. However, the host recognition mechanism of *Siphoviridae*, which recognizes GroP WTA, remains poorly understood. To analyze the host recognition mechanism of our SP phages, we generated phage-resistant *S. pseudintermedius* SP015.

Reduced adsorption of three SP phages was observed in phage-resistant SP015 R1 (Fig 3). Using primers targeting the WTA gene cluster in SP015, we identified a point mutation in *tagN* of R1 that caused a premature stop codon at amino acid 629 (Fig 3a). Complementation of *tagN* using a wild-type allele restored adsorption of SP phages around 50%. Notable, mutation in R1 improved adsorption of ɸSA039 (56.20%), whereas complementation of *tagN* in R1 decreased adsorption of ɸSA039 (31.70%). In SP015 and its mutant derivatives, adsorption did not differ significantly between mutant ɸSA039 (ɸM1, ɸM2, ɸrM1/r-100, and ɸrM1/r-100&102) and wild-type ɸSA039 (Supplemental Figure S2).

**Fig 3.**
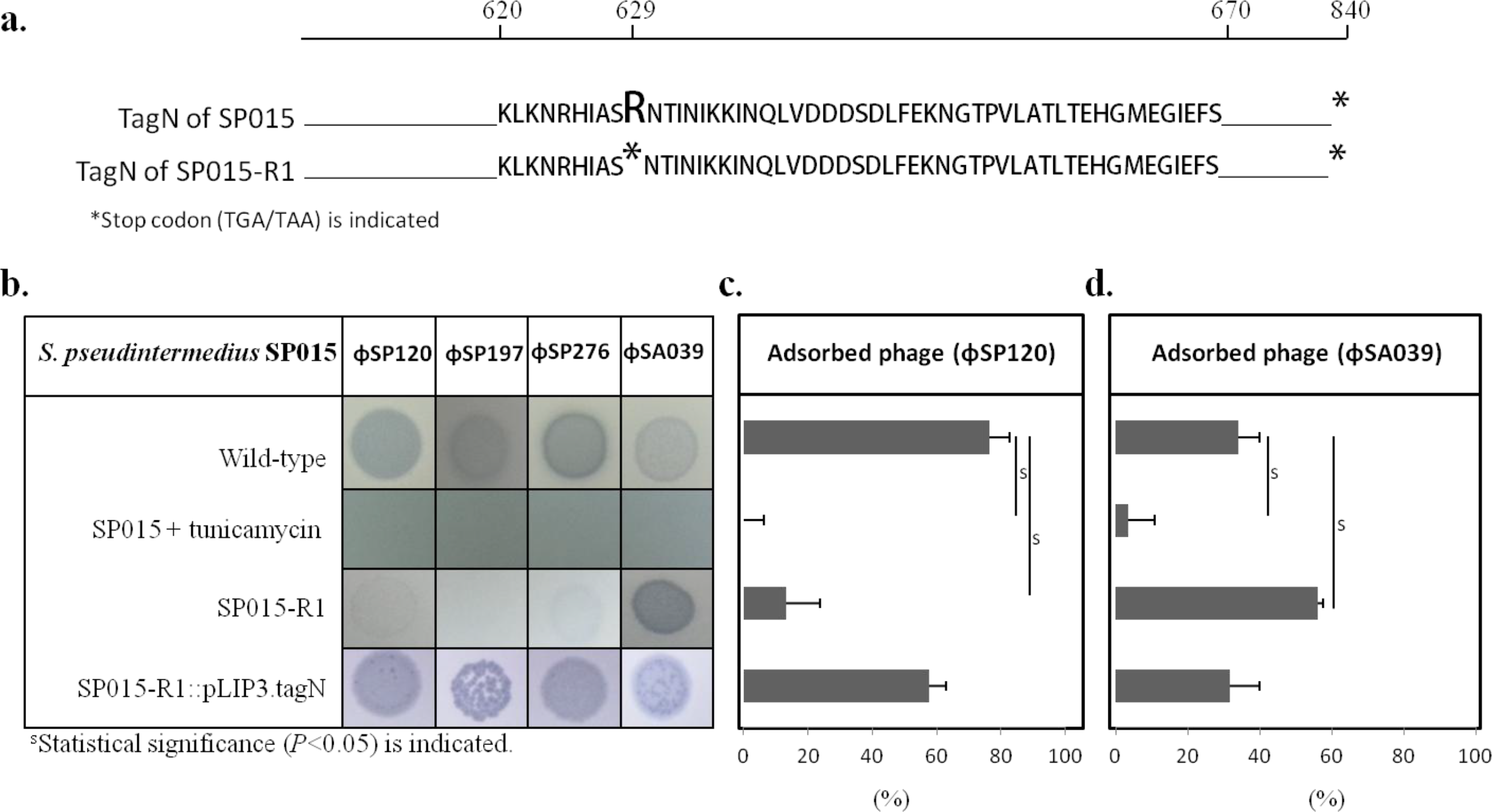
(a) Mutation that causes an early stop codon in *tagN* of SP015-R1. Susceptibility of SP015 wild-type, tunicamycin-induced Sp015 (final concentration 5µg/ml), *tagN* spontaneous mutant (SP015-R1), and TagN-complemented SP015-R1 to all phages, as determined by spot test (b); to phage ɸSP120, as determined by adsorption assay (c); and against phage ɸSA039, as determined by adsorption assay.

## Discussion

### Alteration of RBP enables ɸSA039 to switch its receptor

Although ɸSA039 requires the β-GlcNAc moiety in WTA of *S. aureus*, it can switch its receptor by acquiring spontaneous mutation in its RBP. Alteration of phage receptors is a bacterial strategy used to prevent the initial step of phage infection (Capparelli et al. 2010; Hyman and Abedon 2010; Azam and Tanji 2019a). A previous study showed that spontaneous mutant *S. aureus* lacking the β-GlcNAc moiety in WTA can be easily obtained by co-culturing the bacteria with phage (Azam et al. 2018), suggesting that the emergence of phage-resistant *S. aureus* lacking β-GlcNAc moiety in WTA is possible in a real-world setting. However, in contrast to the situation with antibiotic resistance, ɸSA039 can counter-adapt phage-resistant host by acquiring mutations in ORF100. We confirmed that mutation in ORF100 is key to ɸSA039’s ability to infect TarS-null *S. aureus*. Spontaneous mutants with three point mutations in ORF100 could infect a TarS-null host. Chimeric phages harboring mutant ORF100, produced by homologous recombination, also exhibited similar infectivity. However, a phage with mutated ORF100 could not infect *S. aureus* RN4220, which has TarM (an enzyme responsible for glycosylation of α-GlcNAc of WTA). This residue is likely important for the infection of RN4220 by ɸSA039. Mutation in ORF100 caused the loss of binding ability of ɸSA039 onto RN4220. Our hypothesis was consistent with these observations: ɸM2 and ɸrM1/r-100 could infect TarM-null RN4220 (RN4220ΔTarM), and complementation of TarM in RN4220ΔTarM blocked the infection of the phages.

Subsequent analysis revealed that a point mutation in ORF102 enabled the mutant phage to adsorb onto RN4220. Phage ɸM1, which has mutations in ORF100 and ORF102, was able to bind WTA of RN4220. Our hypothesis is consistent with the ability of a chimeric phage harboring point mutations in ORF100 and ORF102 (ɸrM1/r-100&102) to infect RN4220.

A similar phenomenon has been documented in phages infecting *Escherichia coli*: mutation in *gp38*, which encodes the tail protein of coliphage PP01, enables PP01 to infect *E. coli* O157:H7 lacking the receptor OmpC (Morita et al. 2002). Via such mutations, ɸSA039 and PP01 may acquire the ability to target new receptors other than their original cognate receptors. These indicate that even if the phage-resistant bacteria may appear, new infectious phage will nevertheless be available.

### Staphylococcal kayvirus from family *Myoviridae* is a suitable candidate for phage therapy

Staphylococcal *Myoviridae* phages have a broad host range (Lobocka et al. 2012; Cui et al. 2017). In particular, kayvirus can infect multiple species of *Staphylococcus* (Lobocka et al. 2012). *Staphylococcus* species harbor various glycoepitopes (GlcNAc, GalNAc, or Glc) and two types of WTA backbone (RboP and GroP) (Endl et al. 1983; Brown et al. 2013). Commonly, the infectivity of staphylococcal phages that use glycoepitopes in the WTA as their receptors, e.g. phages from families *Siphoviridae* and *Podoviridae*, depend on the WTA type of the host. For example, *Siphoviridae* phage ɸ187 (a *S. aureus* ST395 specific phage) strictly requires the GalNAc residue in GroP WTA. The phage is not infective toward *S. aureus*, which has RboP WTA (Winstel et al. 2014; Li et al. 2015). The phage can infect coagulase-negative staphylococci (CoNS) including animal-related CoPS, *S. pseudintermedius* (Winstel et al. 2014). The same study also showed that *Siphoviridae* phage ɸ11 and ɸ80α (*S. aureus–*specific phages) can infect *S. aureus* but not *Staphylococcus* species with GroP WTA. Therefore, the host range of phages belong to this group likely depends on the type of WTA.

In this study, we found that ɸSA039 could infect different *Staphylococcus* species with GroP or RboP WTA. The phage exhibited at least moderate infectivity toward *S. pseudintermedius* isolates and strong infection toward CoNS, all of which lack of the β-GlcNAc residue in their WTA and have a GroP backbone. We confirmed that ɸSA039 can utilize GroP-or RboP WTA in *Staphylococcus* bacteria as its receptor. TagN-mediated glycosylation of WTA in *S. pseudintermedius* SP015 likely inhibits the infection of ɸSA039.

Staphylococcal *Siphoviridae* phage often exhibit broad host range, but are specific for bacteria within the same species (Xia et al. 2011; Li et al. 2015). In this study, we demonstrated that SP phages can engage in inter-species infectivity in *Staphylococcus* that possess GroP WTA. Three SP phages (ɸSP120, ɸSP197, and ɸSP276) exhibited strong infectivity toward 19 isolates of *S. pseudintermedius* and 11 CoNS isolates. Unlike other CoNS, *S. saprophyticus* i.e. CCM883 strain has RboP-type WTA (Endl et al. 1983); hence, SP phages cannot infect this species. The absence of non-beneficial genes (toxin, virulence, and antibiotic resistance) in the genomes of ɸSP120, ɸSP197, and ɸSP276 lead us to consider them as potential antimicrobial agents for control of *S. pseudintermedius* infection.

Phage host range is an important criterion when considering a candidate phage for therapeutic application. According to our results, staphylococcal *Myoviridae*, especially those of kayvirus group, are the best candidates in terms of host range; however, to establish efficient treatment, it will be necessary to precisely determine the bacteria responsible for each infection. For example, if one desires to treat *S. pseudintermedius* infection without affecting *S. aureus*, the use of a SP *Siphoviridae* phage represents an alternative. For practical phage therapy, the use of strictly lytic phages that lack non-desirable genes would preferable. Although our newly isolated SP phages are lysogenic, non-desirable genes are absent from their genome; thus, selection of spontaneous mutant phages that are exclusively lytic would be beneficial for practical applications. A recent report described a practical method for screening for spontaneous mutant lytic phages from a lysogenic strain (Gutiérrez et al. 2018), suggesting promising future applications of such phages in therapy.

### Glycosylation of RboP WTA protects *S. aureus* cell from infection by SP phages

Surface carbohydrate moieties are essential for bacterial communication and phage–bacteria and host–pathogen interactions (Weidenmaier et al. 2005; Brown et al. 2013). Most *S. aureus* strains produce RboP WTA substituted with α- and/or β-GlcNAc residues. Many *S. aureus* strains have lost a major genetic barrier against phage infection, the CRISPR-Cas system (Brussow et al. 2004). Consequently, *S. aureus* frequently exchanges genetic material via phage-mediated horizontal gene transfer (HGT) events (Xia and Wolz 2014; Li et al. 2015). The difference in WTA structure determines efficient HGT among *Staphylococcus* bacteria. Staphylococcal *Siphoviridae* mediates HGT among *Staphylococcus* (Xia and Wolz 2014). Host recognition of phages from this family depends on WTA structure (Azam and Tanji 2019b). For example, *Siphoviridae* phages capable of infecting *S. aureus* (SA *Siphoviridae*), such as ɸ11 and ɸ80A, can recognize GlcNAc of RboP WTA, but cannot recognize GroP WTA of CoNS (see illustration in Fig 4). On the other hand, *Siphoviridae* that infect other *Staphylococcus* species have rarely been reported.

**Fig 4.**
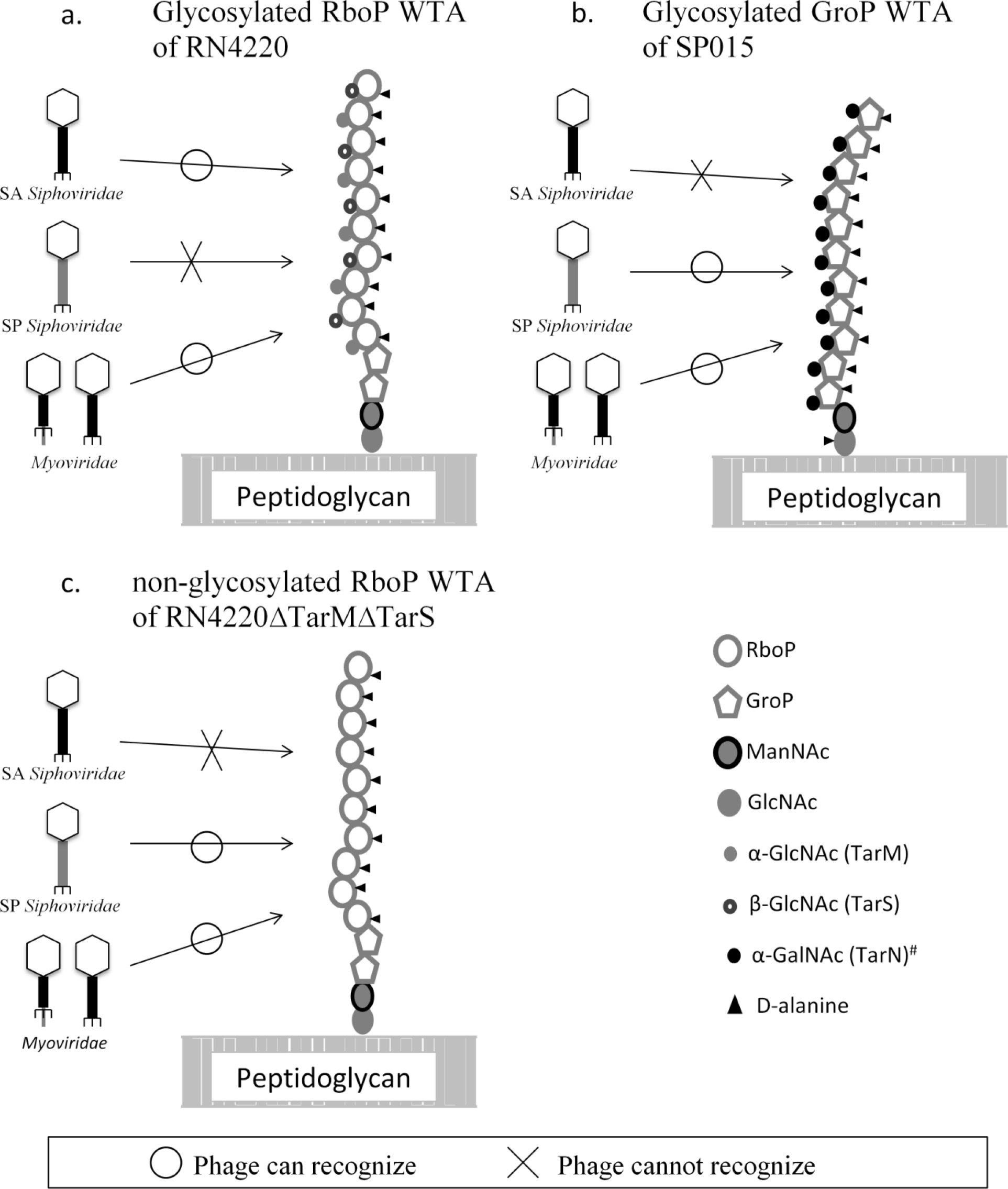
Mechanism of recognition of WTA by staphylococcal phages: (a) SA *Siphoviridae* and *Myoviridae* recognize glycosylated RboP WTA; (b) SP *Siphoviridae* and *Myoviridae* recognize glycosylated GroP WTA; and (c) SP *Siphoviridae* and *Myoviridae* recognize non-glycosylated RboP WTA. *Glycosylation by TagN is based on a report in the *S. aureus* ST395 lineage (Winstel et al. 2014).

Within *S. aureus* species, the ST395 lineage harbors a unique WTA containing 1,3-glycerol-phosphate (GroP) modified with α-GalNAc and D-alanine (Winstel et al. 2013; Winstel et al. 2014). The WTA structure of the ST395 lineage resembles that of CoNS and *S. pseudintermedius*. *Siphoviridae* phages capable of infecting PS187 (a representative strain of the ST395 lineage) can also infect CoNS and other species with similar WTA structure (Winstel et al. 2014). In this study, we characterized *Siphoviridae* phages capable of infecting *S. pseudintermedius* (termed SP *Siphoviridae*). We demonstrated that the *S. pseudintermedius* strains used in our study have GroP WTA. In a representative strain (SP015), we observed that our SP *Siphoviridae* phages required glycosylated GroP WTA. A nonsense mutation resulting in deletion of the C-terminus of *tagN*, encoding glycosyl transferase for GroP WTA, caused a significant decrease in adsorption of SP phages onto SP015 but improved adsorption of ɸSA039. Adsorption of phages was not observed in the present of tunicamycin (an antibiotic which inhibits WTA synthesis), suggesting that SP phages and ɸSA039 required WTA for infection but glycosylation by TagN was dispensable for ɸSA039.

Interestingly, we showed that SP phages could recognize *S. aureus* harboring non-glycosylated WTA (due to lack of TarM and TarS). Although *S. aureus* lacking both TarM and TarS is not likely to be present in nature, our finding indicates that glycosylation of RboP WTA of *S. aureus* is a strategy used by the bacteria to protect themselves against infection by SP *Siphoviridae*, thereby limiting the HGT across different species with different WTA backbones. Similar strategies have been documented in other reports (Li et al. 2015) showing that TarM protects *S. aureus* against the lytic activity of *Podoviridae*. Staphylococcal *Podoviridae* require precise WTA glycosylation pattern for infection. TarM-mediated α-GlcNAc glycosylation in RboP WTA prevents infection of *Podoviridae* while TarS-mediated β-GlcNAc glycosylation is important for *S. aureus* susceptibility to *Podoviridae*. Our findings reveal a novel strategy by which *S. aureus* protects itself against infection by phages capable of specifically infecting phylogenetically distant species. These finding provide novel insight into biology of staphylococcal phage. We suggested that host recognition mechanism of staphylococcal phage is likely more complex than our current understanding.

## Acknowledgements

We thank Professor Kenji Kurokawa at the Faculty of Pharmaceutical Science of Nagasaki International University for providing us with the deletion mutant of RN4220.

## Compliance with ethical standards

Conflict of interest: The authors declare that they have no competing interests.

Ethical approval: This article does not contain any studies with human participants or animal performed by any of the authors.

---

**Supplemental Table S1.**
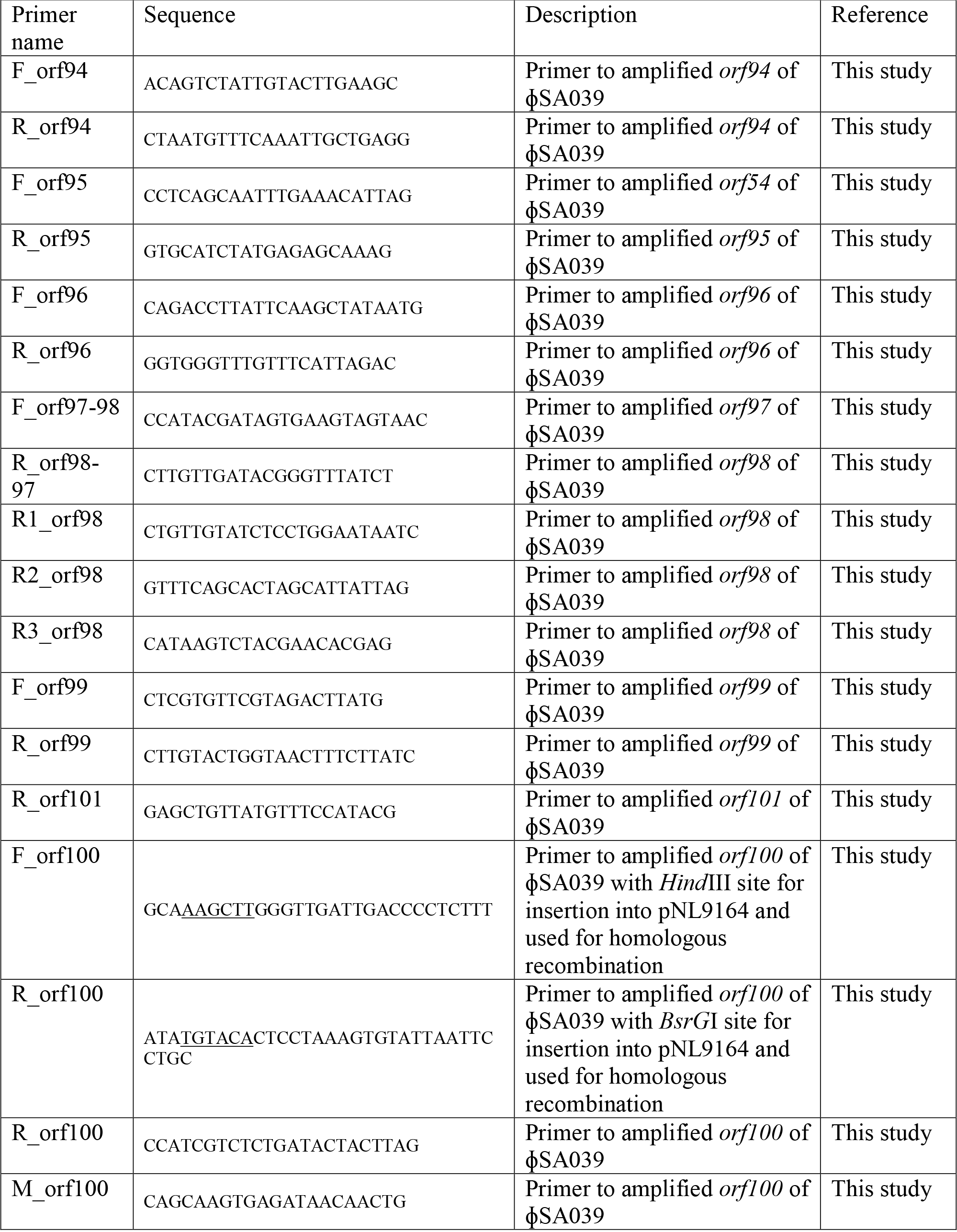
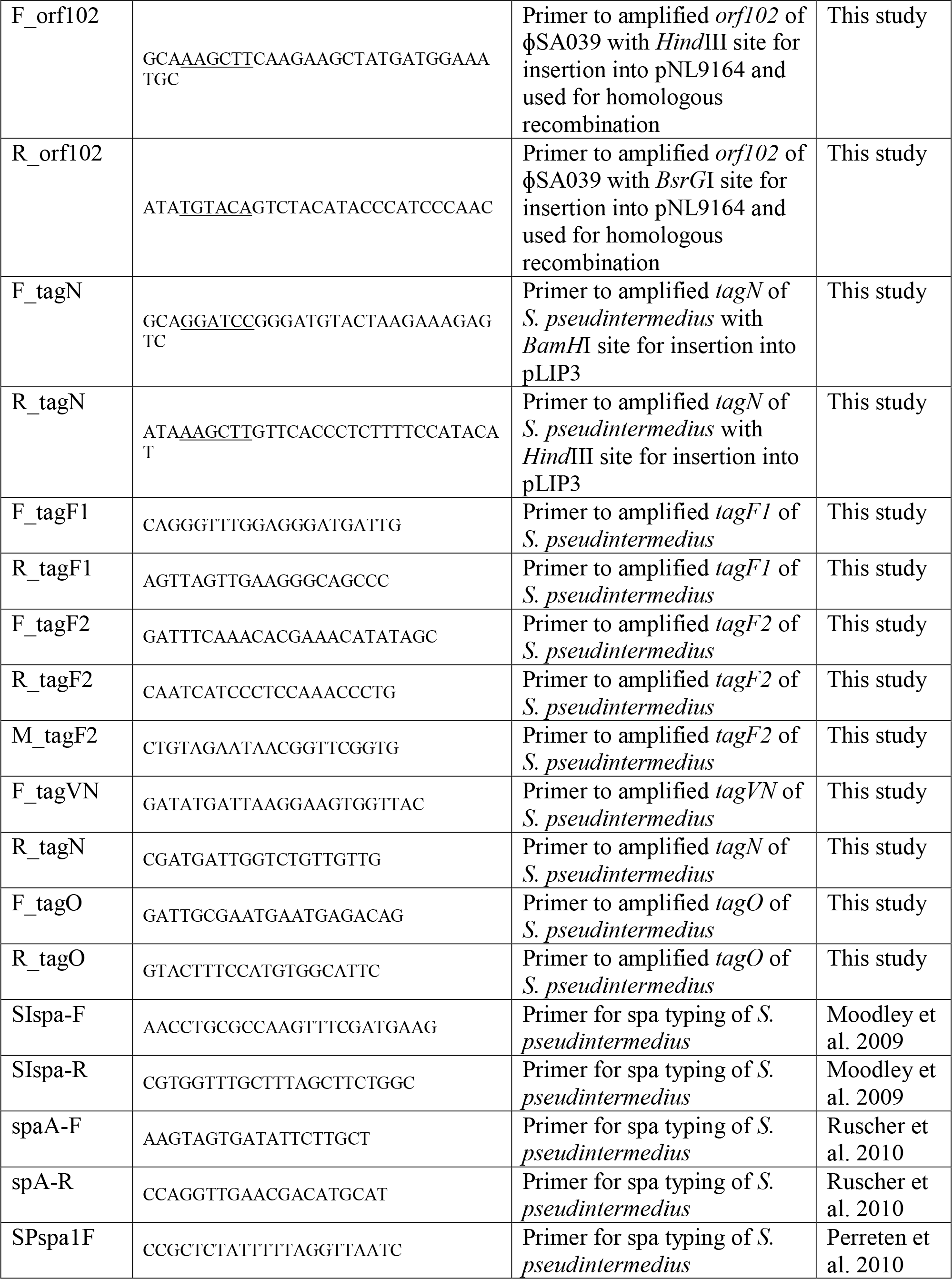
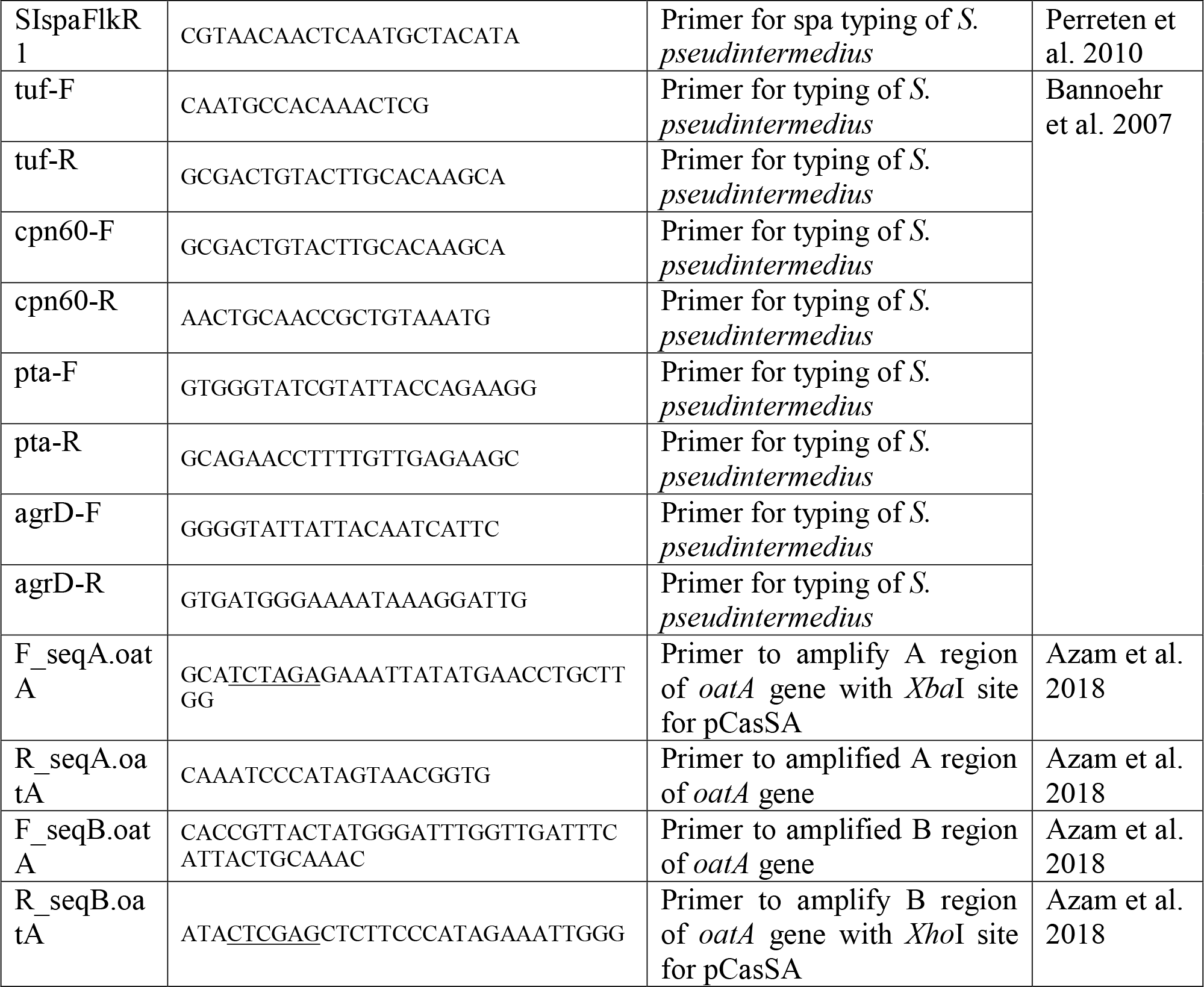
Primers used in this study.

**Supplemental Figure S2.**
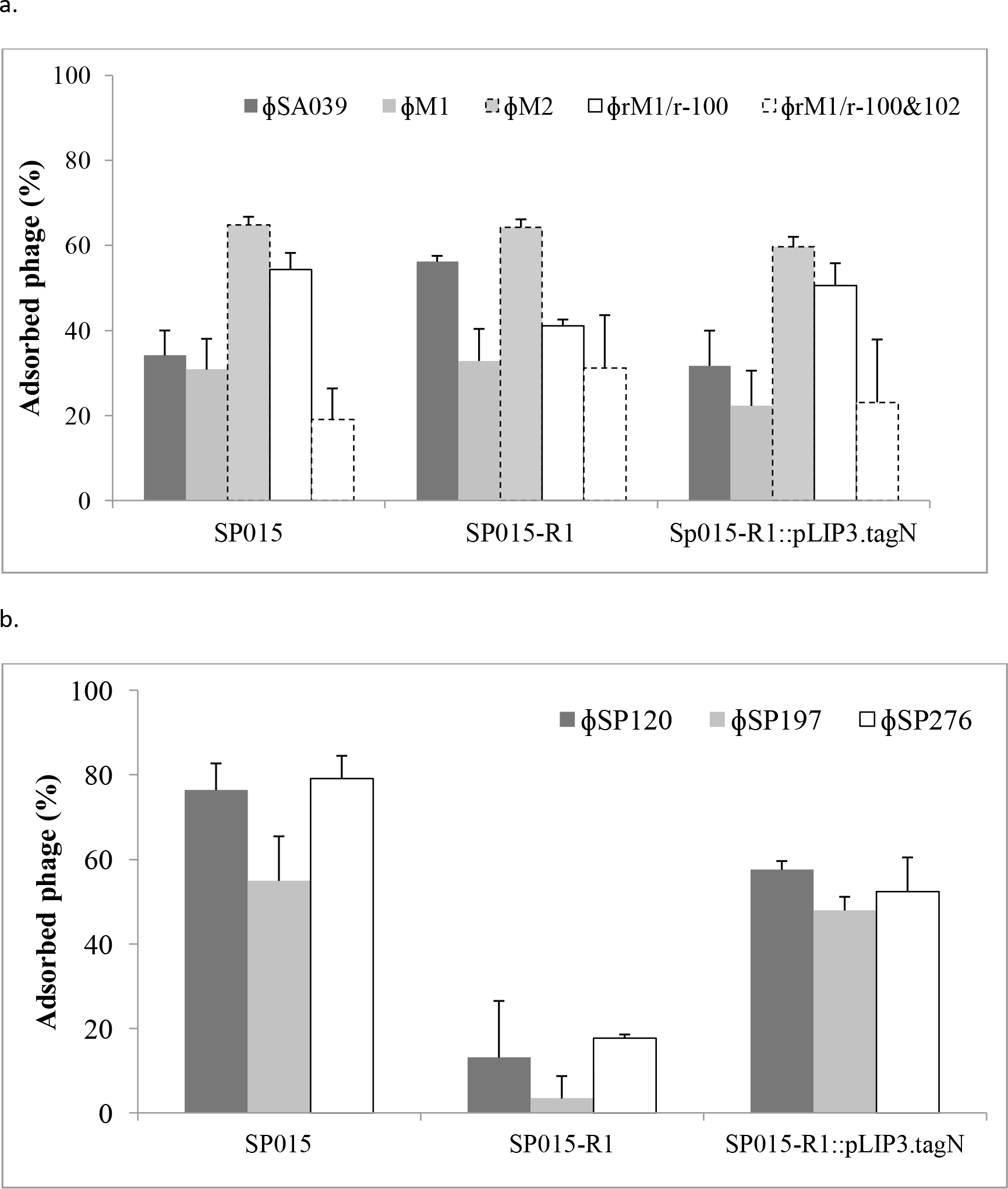

**Supplemental Table S3.**
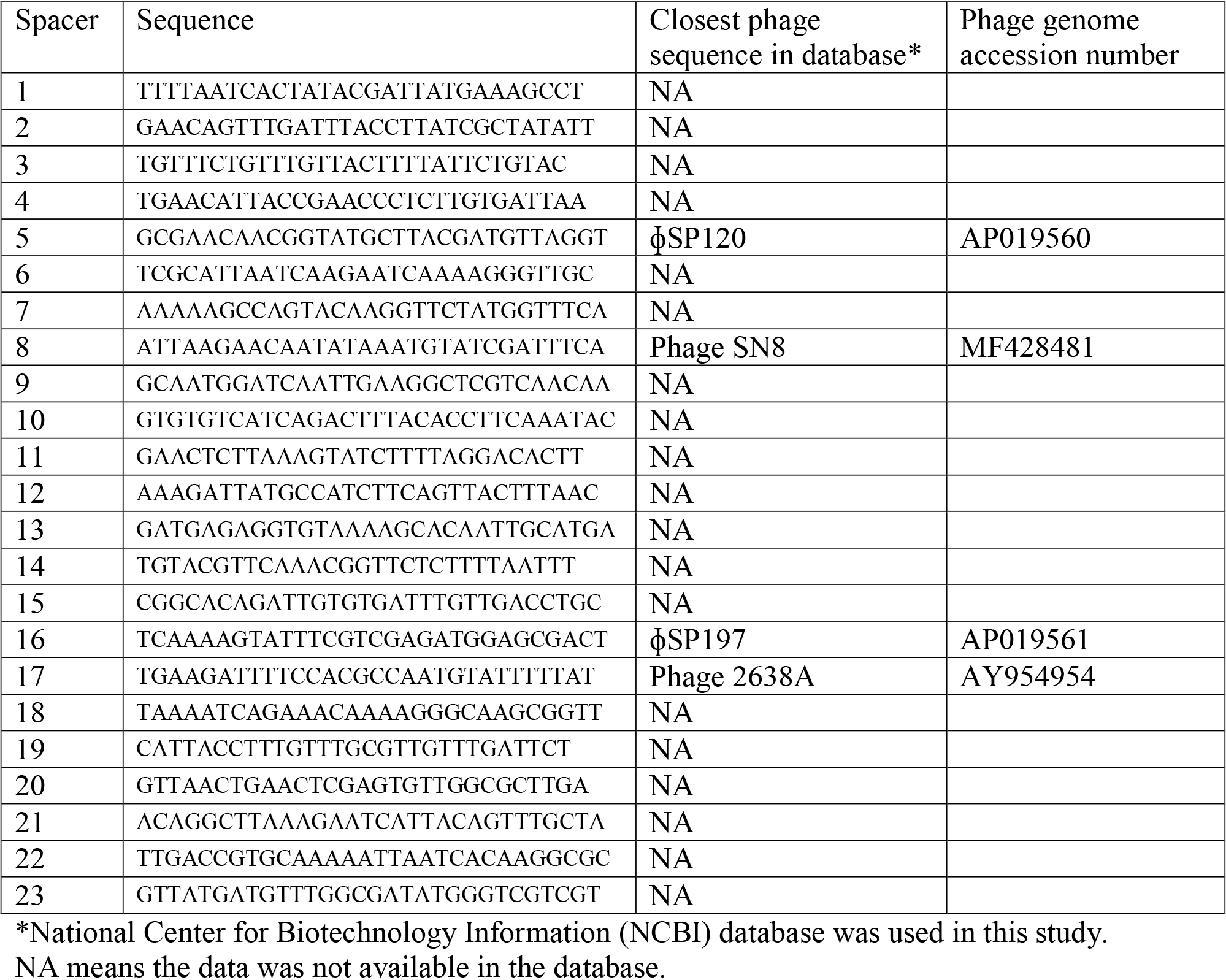
Spacers present in CRISPR of *S. pseudintermedius* SP079

